# Identification of new viruses specific to the honey bee mite *Varroa destructor*

**DOI:** 10.1101/610170

**Authors:** Salvador Herrero, Sandra Coll, Rosa M. González-Martínez, Stefano Parenti, Anabel Millán-Leiva, Joel González-Cabrera

**Affiliations:** ERI BIOTECMED. Department of Genetics. Universitat de València, Valencia, Spain

**Keywords:** iflavirus, picornavirus, insobevirus, qPCR, +ssRNA virus

## Abstract

Large-scale colony losses among managed Western honey bees have become a serious threat to the beekeeping industry in the last decade. There are multiple factors contributing to these losses but the impact of *Varroa destructor* parasitism is by far the most important, along with the contribution of some pathogenic viruses vectored by the mite. So far, more than 20 viruses have been identified infecting the honey bee, most of them RNA viruses. They may be maintained either as covert infections or causing severe symptomatic infections, compromising the viability of the colony. *In silico* analysis of available transcriptomic data obtained from mites collected in the USA and Europe as well as additional investigation with new samples collected locally allowed the description of three novel RNA viruses. Our results showed that these viruses were widespread among samples and that they were present in the mites and in the bees but with differences in the relative abundance and prevalence. However, we have obtained strong evidence showing that these three viruses were able to replicate in the mite, but not in the bee, suggesting that they are selectively infecting the mite. To our knowledge, this is the first demonstration of Varroa-specific viruses, which open the door to future applications that might help controlling the mite through biological control approaches.

## Introduction

The ectoparasitic mite *Varroa destructor* Anderson and Trueman (Arachnida: Acari: Varroidae) is considered a major driver for the seasonal losses of managed Western honey bee (*Apis mellifera* L.) colonies reported worldwide (Rosenkranz et al. 2010). There are multiple factors contributing to these losses, but the impact of *V. destructor* is by far the most important in many cases (Roberts et al. 2017). The damage caused by the mite is a result of the combined effect of direct feeding on the fat body of immature and adult bees (Ramsey et al. 2019) and the transmission of an extensive set of viruses that infect and debilitate them (McMenamin and Genersch 2015). This resulted in reduced immune response, low tolerance to pesticides, impaired mobility or flying capacity, reduced weight, morphological deformities, paralysis, etcetera (De Jong et al. 1982; Yang and Cox-Foster 2007; Yang and Cox-Foster 2005).

So far, more than 20 viruses have been identified in the honey bees (McMenamin and Genersch 2015), most of them ssRNA viruses. Among them, the Acute bee paralysis virus (ABPV), Deformed wing virus (DWV), Israel acute paralysis virus (IAPV), Kashmir bee virus (KBV), Sacbrood virus (SBV), Slow bee paralysis virus (SBPV), Chronic bee paralysis virus (CBPV), Lake Sinai virus (LSV), Kakugo virus, *Varroa destructor* virus-1 and Black queen cell virus (BQCV) (Boecking and Genersch 2008; Di Prisco et al. 2011; McMenamin and Genersch 2015; Santillán-Galicia et al. 2014). However, in absence of *V. destructor* parasitism, many of these viruses have been found infecting seemingly healthy honey bee colonies, in what are considered as covert infections, that do not cause any detectable clinical impact on them (McMenamin and Genersch 2015). After the host shift of *V. destructor* from *Apis cerana* L. to *A. mellifera* L., the prevalence and pathogenicity of some of the aforementioned viruses have increased significantly (Rosenkranz et al. 2010). This combination of viruses’ pathogenicity and the high vectoring capacity of *V. destructor* is considered for many as one of the main causes of the Colony Collapse Disorder (CCD), responsible for very high losses of colonies in Europe and the USA (Potts et al. 2010; Vanengelsdorp et al. 2009).

Thus, it is well documented the ‘relation’ of *V. destructor* with bee pathogenic viruses but there is very little information available about viruses specific (or pathogenic) for the mite itself. There are no evidences of bee pathogenic viruses causing any impact on *V. destructor* physiology, even with high loads of some of them recorded in mite virome. There are even viruses discovered in the mite whilst showing no evidence of pathogenicity for it (i.e. VdV-1) (Ongus et al. 2004). Recent research has identified two novel viruses present in *V. destructor* but not detected in the bees. However, the information available does not show clear evidences of the possible differential specificity of these viruses for *V. destructor* or the honey bee (Levin et al. 2016).

In this paper, we report the identification and phylogenetic characterization of three new viruses in the virome of *V. destructor*. We show strong evidence indicating that these viruses can replicate in the mite but not in the bee, suggesting that they are selectively infective for the mite without affecting the replication of DWV.

## Material and Methods

### Mite samples

Mites used in this study were collected directly from the brood of honey bee combs, from hives located nearby Valencia, Spain. Capped cells were opened with a pair of tweezers to extract immature bees and the mites parasitizing them. Bees and mites extracted from the same cell were put together in a microcentrifuge tube, snapped frozen in liquid nitrogen and stored at −80 °C until used for qPCR analysis.

### Virus discovery and genome sequence analysis

Viral sequences were identified in two *V. destructor* transcriptomes assembled from data downloaded from the SRA (https://www.ncbi.nlm.nih.gov/sra). The mites used to generate these data were collected in the USA (Accession SRS353274) and in the UK (Accession PRJNA531374). A set of invertebrate picornaviral sequences from different families was used as query in Blastx search against the different transcriptomes. Those sequences producing significant hits were manually curated to remove redundant sequences. Sequences with similarity restricted to functional domains like zinc finger domain, helicase domain, and RNA binding domain were removed since they can be in viral as well as in non-viral proteins, and possibly representing non-viral sequences.

The genomic structure, gene content, and location of the conserved motifs for the non-structural proteins (helicase, protease and RNA-dependent RNA polymerase), were obtained by comparison with their closest viral species.

### Phylogenetic analysis

RNA-dependent RNA polymerase was selected to establish the phylogenetic relationship of viruses described in this study with the closest viruses and representative members of nearest viral families, retrieved from NCBI GenBank (www.ncbi.nlm.nih.gov/genbank).

Protein sequences comprising putative conserved domains for +ssRNA viruses were aligned using MAFFT version 7.409 software employing the E-INS-i algorithm (Standley and Katoh 2013). Alignments were examined by eye and subsequently, ambiguously aligned regions were removed using TrimAl program (Capella-Gutiérrez et al. 2009). The most appropriate evolutionary model (best-of-fit model) according with corrected Akaike Information Criterion (AICc) was calculated using Small Modelling Selection software (Longueville et al. 2017). Phylogenetic trees were inferred by maximum likelihood approach implemented in PhyML version 3.1 (Guindon et al. 2010), with the LG substitution model, empirical amino acid frequencies, and a four-category gamma distribution of rate, and using Subtree Pruning and Regrafting (SPR) branch-swapping. Branch support was assessed using the approximate likelihood ratio test (aLRT) with the Shimodaira–Hasegawa-like procedure as implemented in PhyML. Branches with <0.6 aLRT support were collapsed using TreeGraph software (Stöver and Müller 2010).

### Virus detection and quantification

Presence and abundance of RNA viruses in honey bee and *V. destructor* samples were determined by detection of viral RNA genomes using reverse transcription quantitative real-time polymerase-chain reaction (RT-qPCR). For this purpose, total RNA was isolated from individual *V. destructor* and honey bees using Trizol reagent (Sigma Aldrich). For the mites, each individual was homogenized with a plastic pestle in microcentrifuge tubes containing 100 μl of Trizol in two cycles of freeze and thaw in liquid nitrogen. After that, 100 μl of Trizol were added to the mixture and the RNA extraction continued following the manufacturer’s protocol. Total RNA from honey bees was isolated from the abdomen of individual bees after homogenization in 200 μl of Trizol reagent using the TissueLyser LT (Qiagen N. V.) for 3 min. For cDNA synthesis, 300 ng of each RNA was reverse transcribed to cDNA using random hexamers and oligo(dT) primers and following the instructions provided in the PrimeScript RT Reagent Kit (Perfect Real Time from Takara Bio Inc., Otsu Shiga, Japan). RT-qPCR was carried out in a StepOnePlus Real-Time PCR System (Applied Biosystems, Foster City, CA). All reactions were performed using 5× HOT FIREpol EvaGreen qPCR Mix Plus (ROX) from Solis BioDyne (Tartu, Estonia) in a total reaction volume of 20 μl.

Forward and reverse primers used in the study were either described elsewhere or designed *de novo* for this study using the Prime3Plus software (Nijveen et al. 2007) (Table 1). Primers efficiency were calculated for all primer pairs used in RT-qPCR (Table 1). The ribosomal 18S gene (Campbell et al. 2016) and the ApiDorsal gene (Di Prisco et al. 2016) were used as endogenous control of the RNA concentration for the varroa and bee samples, respectively. As a preliminary outcome of the RT-qPCR, the identity of amplified fragments was confirmed by Sanger sequencing. For relative viral load quantification in each of the samples, the following equation was used, Relative abundance = E (primers virus)^−Ct (virus)^ / E (primers endogenous)^−Ct (endogenous)^, were E, refers to the efficiency of each pair of primers and Ct the values of the Threshold Cycle obtained in the qPCRs. The limit of detection for each virus was established based on the average Ct values obtained for the negative controls (total RNA without cDNA synthesis). The Ct values equal or greater to the average Cts for the controls were considered free of the studied virus and reported as non-detected (ND).

Semi-quantitative RT-PCR was used to detect the negative RNA strands of the VdV-5, VdV-3 and VdIV-2 in varroa and bees as previously described (Jakubowska et al. 2015; Llopis-Giménez et al. 2017). Briefly, tagged primer was used for the specific synthesis of cDNA due to the occurrence of self-priming, which is often observed for RNA viruses. Total RNA was extracted as described above from a pool of about 10 varroa mites, or 10 bee heads or abdomens. For this, 500 ng of RNA were used for cDNA synthesis using a tagged specific primer (Table S1). cDNA synthesis was performed using the PrimeScript RT reagent kit from Takara Bio Inc (Otsu Shiga, Japan) at 42 °C for 30 min, following the manufacturer’s protocol. Three microliters of cDNA were used for subsequent PCR reactions using the tag region as a forward primer, and specific reverse primers (Table S1). PCR was performed using the following conditions: 95 °C for 5 min, followed by 35 cycles of annealing at 52 °C for 30 seconds, and elongation at 72 °C for 1 min using DreamTaq DNA polymerase (Thermo Fischer Scientific, Waltham, MA, USA). Presence of viral positive strand was confirmed in each of the samples used for the negative strand detection by RT-qPCR as described above.

## Results and discussion

### New viruses associated to *V. destructor*

The presence of virus-like sequences associated with *V. destructor* was investigated via data mining using transcriptomic data available in public databases. These analyses showed that among other viruses, typically associated with the vectoring capacity of *V. destructor* (data not shown), the data contained sequences belonging to three different +ssRNA viruses featuring conserved regions and motifs suggesting that they were novel. Detailed analyses of the selected sequences revealed the absence of mutations or indels generating premature stop codons, which supported the idea that all three sequences belong to viruses actively infecting *V. destructor* (Fig. 1–4). Two of the viruses showed significant similarity with a virus recently described that was named *Varroa destructor* virus 3 (VdV-3) (Accession KX578272.1) (Levin et al. 2016). Here, we named that virus VdV-3 ISR to differentiate it from a closely related variant found in our data. This variant showed 90.8 % overall genome identity with VdV-3 ISR. Thus, we consider it a new variant of the same species and we named it VdV-3 USA. The other virus similar to VdV-3 ISR showed an overall identity of only 75.0 %. Therefore, we consider it a new species and it was named *Varroa destructor* virus 5 (VdV-5) (Fig. 1A). Despite differences in the amino acid sequence, the genome of the two viruses described here were very similar in structure. Two coding Open Reading Frames (ORF) were predicted for both viral genomes, ORF1 with 817 and 822 amino acids in VdV-3 USA and VdV-5, respectively and ORF2 with 493 amino acids. In ORF1 we identified conserved motifs generally found in closely related species: a transmembrane domain (TM), a trypsin-like serine protease (Pro) and a viral genome-linked protein (VPg). On the other hand, ORF2 contained only conserved motifs for the RNA-dependent RNA polymerase (RdRp) (Fig. 1A).

**Figure 1.**
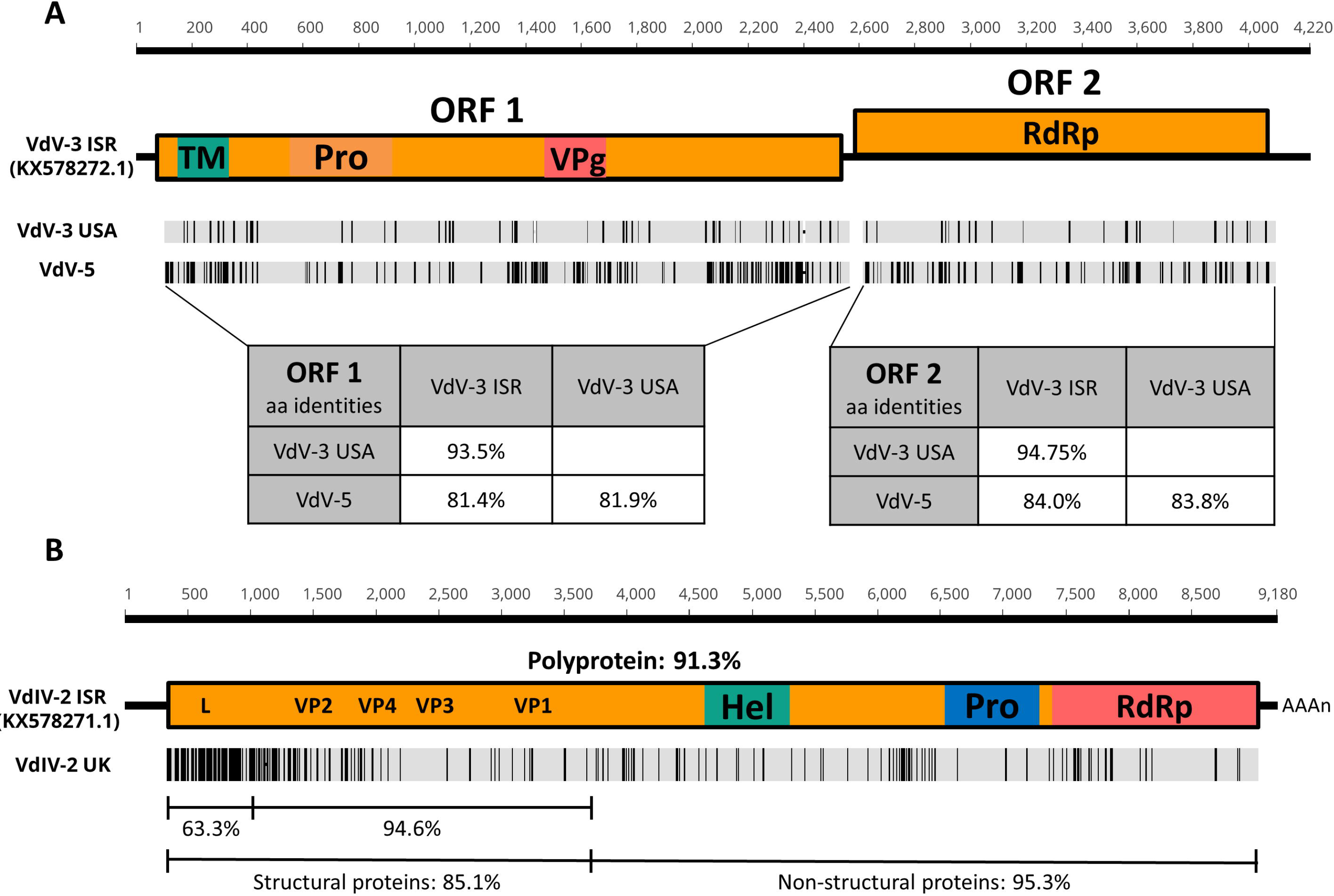
Schematic representation of the new viruses. Schematic representation of the genome organization for the *Varroa destructor* viruses described on this project, compared to the closest previous described VdV-3 (**1A**) and VdIV-2 (**1B**) (accessions KX578272.1 and KX578271.1, respectively). Grey boxes represent ORF and vertical black lines indicate amino acid variation. Percentage values show amino acid sequence identities. Predicted ORFs are shown as boxes. Conserved domains on the amino acid sequence are represented by letters (RdRp: RNA-dependent RNA polymerase; Pro: Protease; Hel: RNA Helicase; VP1 to VP4: Capsid proteins 1 to 4; L: Leader protein; TM: Transmembrane domain; VPg: Viral genome-linked protein). Percentage of aa identities are provided for the different domains of the polyprotein as well as for the different ORFs.

The phylogenetic analysis of the new viruses using the RdRp sequence showed that, given the sequence similarity, they clustered together with VdV-3 ISR in a branch containing the other unassigned invertebrate-derived viruses closely related to plant +ssRNA viruses from the *Solemoviridae* (Sobemovirus, and Polemovirus) and *Luteoviridae* (Enamovirus) families (Fig. 2). They are also closely related with Barnaviruses, described in fungi. Members of this group have a similar genomic structure with two ORFs spanning 3 to 4.2 kb. This group includes viruses and viral sequences described in other acarine species like the deer tick (*Ixodes scapularis*) and the American dog tick (*Dermacentor variabilis*) (Harvey et al. 2019; Levin et al. 2016; Levin et al. 2019; Pettersson et al. 2017; Sadeghi et al. 2018; Shi et al. 2016; Webster et al. 2016) and not much information about their incidence and replication in a specific host has been reported. Nevertheless, given the mentioned similarities and the replicative activity of these viruses (see below), these *V. destructor* associated viruses could define a new family of +ssRNA viruses infecting invertebrates. Thus, we propose to name this new family as “*Insobeviridae*”.

**Figure 2.**
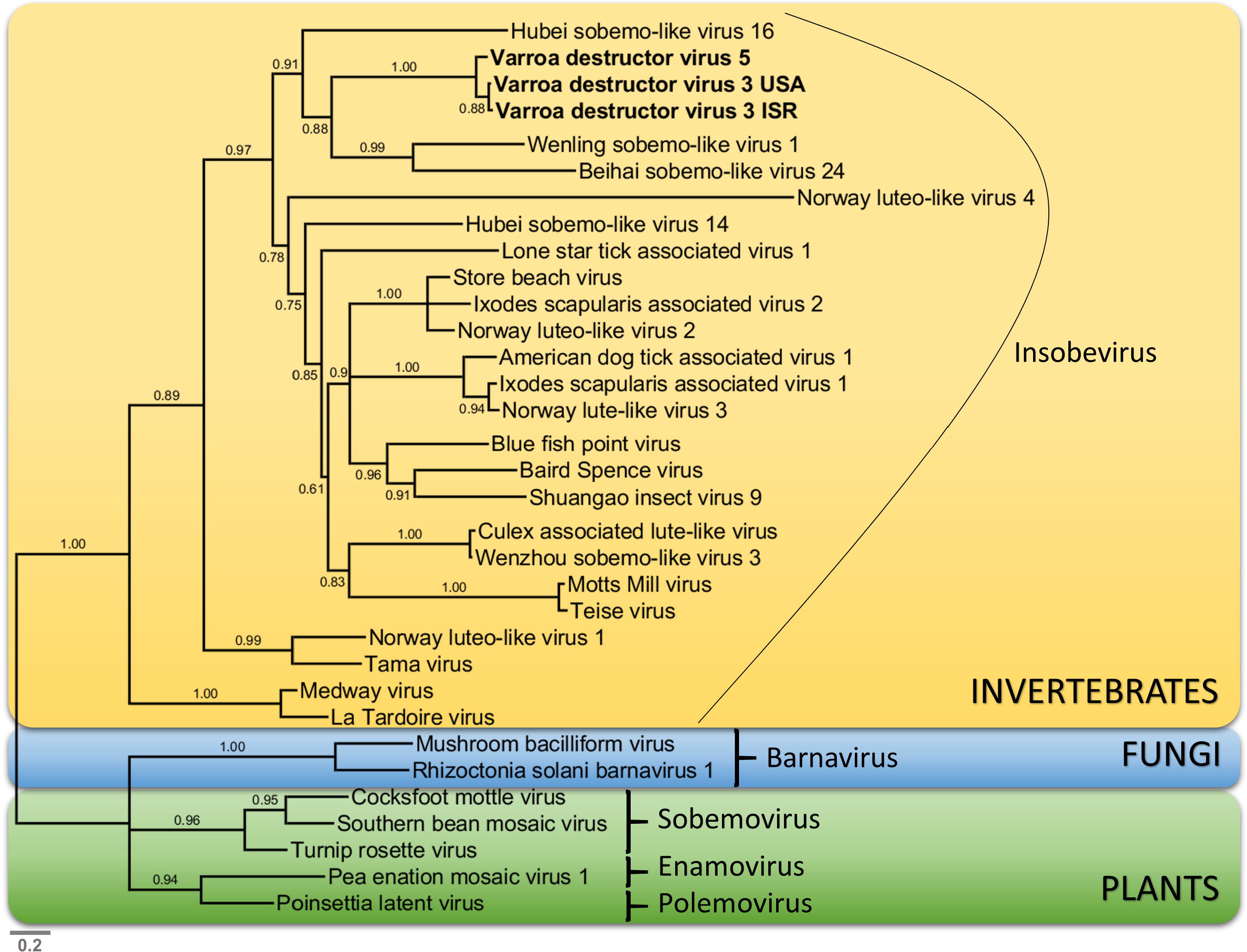
Phylogenetic analysis of the new Insobeviruses and its relationship with other RNA viruses. Maximum-likelihood phylogenetic tree of the putative RdRp amino acid sequences of Varroa destructor virus 5 and 3, and closely related viruses. Values on blanches indicates the aLRT support. Branches with aLRT values <◻0.6 have been collapsed. The grey scale bar represents 0.2 amino acid substitution per site. Sequence accession identifiers are provided on Supplementary Table S2

The third viral sequence showed high similarity to the iflavirus *Varroa destructor* virus 2 described earlier (VdIV-2) (Levin et al. 2016), sharing 83.4 % of genome identity between them (Fig. 1B, S1). Due to the high similarity with the VdV-2, we resolve that they were genotypes/isolates from the same species, hence we named as Varroa destructor iflavirus 2 UK (VdIV-2 UK), to discriminate it from the previous described virus which we named as Varroa destructor iflavirus 2 ISR (VdIV-2 ISR).

The genome of the VdIV-2 UK contains a single large ORF, that translates into 2996 amino acids polyprotein, displaying the characteristic *Iflaviridae* structure organisation (Valles et al. 2017) (Fig 1B). Conserved motifs for the Non-structural proteins, RdRp, 3C-like cysteine protease and RNA helicase, were detected in the C-terminal region of the polyprotein. Capsid proteins were identified based on their homology to those from other members of the family. Polyprotein sequence for the two viruses showed a 91.3 % similarity, nonetheless, most amino acid variations were concentrated on the Structural-proteins coding region, with half of the differences (131 out of 262 total aa changes) on the tentative leader protein (L) sequence, at the polyprotein N-terminus (Fig. 1B).

### Viral incidence in field samples

To validate the actual presence of these viruses in the wild, we designed, synthesized and tested primer pairs specific for each of the three viruses described above (See Material and Methods). The detection was carried out in both, the *V. destructor* individuals parasitizing immature bees (Fig. 3A) and also in these very same bees collected from the same capped cell (Fig. 3B). The results showed that the three viruses were detected in the mites as well as in the bees although with differences in the relative abundance and prevalence. The virus VdV-5 was present in all mites tested while it was only detected in 57 % of individual bees. Virus VdV-3 USA showed the lowest prevalence with detections in 36 % and 43 % of the analysed mites and bees, respectively. On the other hand, the iflavirus, VdIV-2 UK, was found in all tested mites and in most of the bees (86 %) (Fig. 3). It is interesting to point out that in about 30 % of the samples it was possible to detect the three viruses in a single individual (35 % in mites and 29 % in bees). The high incidence of these viruses in our samples (from Spain) and the presence of closely related variants in samples collected in Israel (Levin et al. 2016), the UK and the USA (transcriptomic data used in this study) is a clear indication of their widespread distribution in the populations and that they are evolving, possibly to gain better adaptation to different mite populations and environmental conditions. Whether these new viruses influence the mite or bee performances or not, remain to be elucidated in further studies. However, an important step to study its relevance in the varroa-bee system is to determine their replication and host specificity.

**Figure 3.**
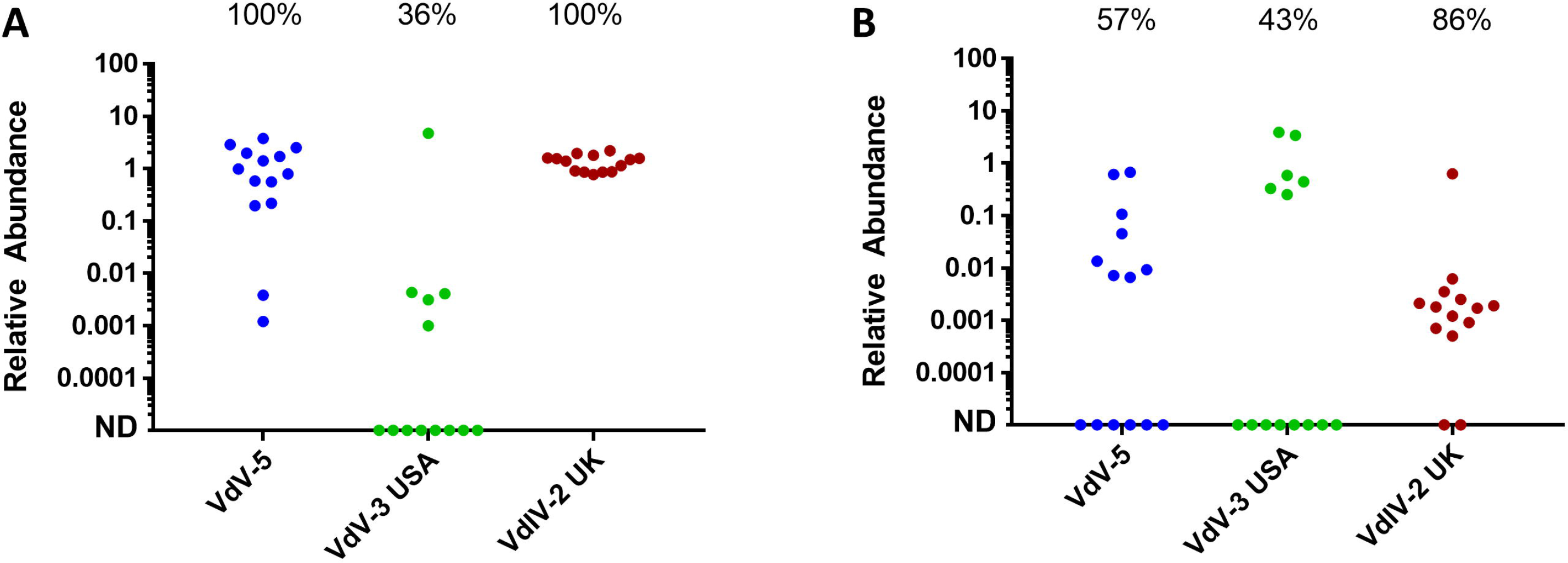
Incidence and relative abundance in varroa and bee individuals. Relative abundance of the viral RNA in Varroa mite (A) parasitizing immature bees (B) obtained by RT-qPCR and normalized according to the expression of their respective reference gen (*18S* for Varroa and *Apidorsal* for the bees). Each spot represents an individual mite or bee. ND (non-detected) is used for those samples scoring under the detection threshold. The percentage of positive individuals is reported in each sample.

### Viral replication and specificity

A +ssRNA virus produces an intermediate negative-strand RNA when it replicates. Thus, the detection of negative-strand viral RNA is indicative of viral replication and supports its viral identity. We designed specific sets of primers to detect the negative strand by PCR, to confirm the viral origin of the sequences as well as to determine their host specificity. The results showed that although it was possible to detect viral genomes in the bee heads and/or abdomens (Fig 4. Upper panels), the negative strand for VdV-5, VdV-3, and VdIV-2 was only detected in the mites (Fig 4), indicating that the sequences described here belong to active viruses and that they replicate exclusively in the mites. It seems that their presence in the bees was probably due to the typical exchange of fluids between the bee and the mite parasitizing it.

**Figure 4.**
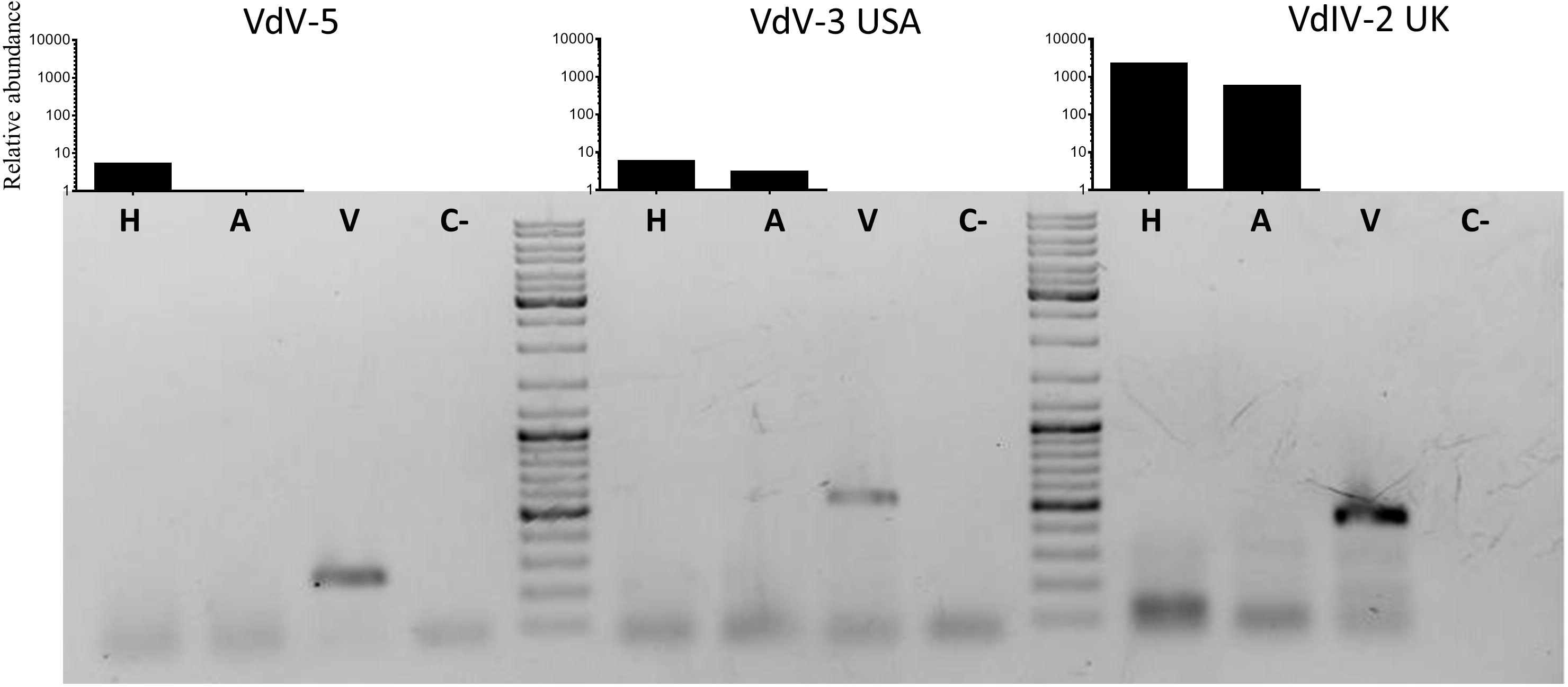
Detection of viral negative strand of the new viruses. Viral negative strand intermediate was detected by specific RT-PCR and agarose electrophoresis. Specific amplification of the three viruses was carried out in the bee heads (**H**) and abdomens (**A**) and the whole varroa (**V**). **C−** represents the negative control. The relative abundance of the viral positive strand was estimated for the same type of samples (upper bar graphs) by RT-qPCR and normalized according to the relative abundance of the bee-specific gene *Apidorsal*.

In addition to the lack of viral negative strand in the bee samples, there are other indirect evidences supporting the specific replication of these new viruses in mite tissues. For example, it is possible to estimate the abundance of mite contents in the parasitized bees from the amplification of the *V. destructor* endogenous gene Vd*18S* in bee samples. We have observed a positive correlation (Pearson r = 0.44; P-value = 0.019) between the viral abundance in the bees and the amount of mite content (Fig 5A). However, the comparison of viral loads in pairs bee/mite did not reveal a correlation of relative abundance of each of the viruses between the mite and its parasitized bee (P-value>0.05 for all the viruses) (Fig. 5B). In fact, it was remarkable that, in some cases, the mite had a very high virus load while it was not possible to detect the same virus in the bee (Fig. 5B). Accordingly, we hypothesised that the presence of viral sequences in the bees is just circumstantial and does not reflect viral load on the mite.

**Figure 5.**
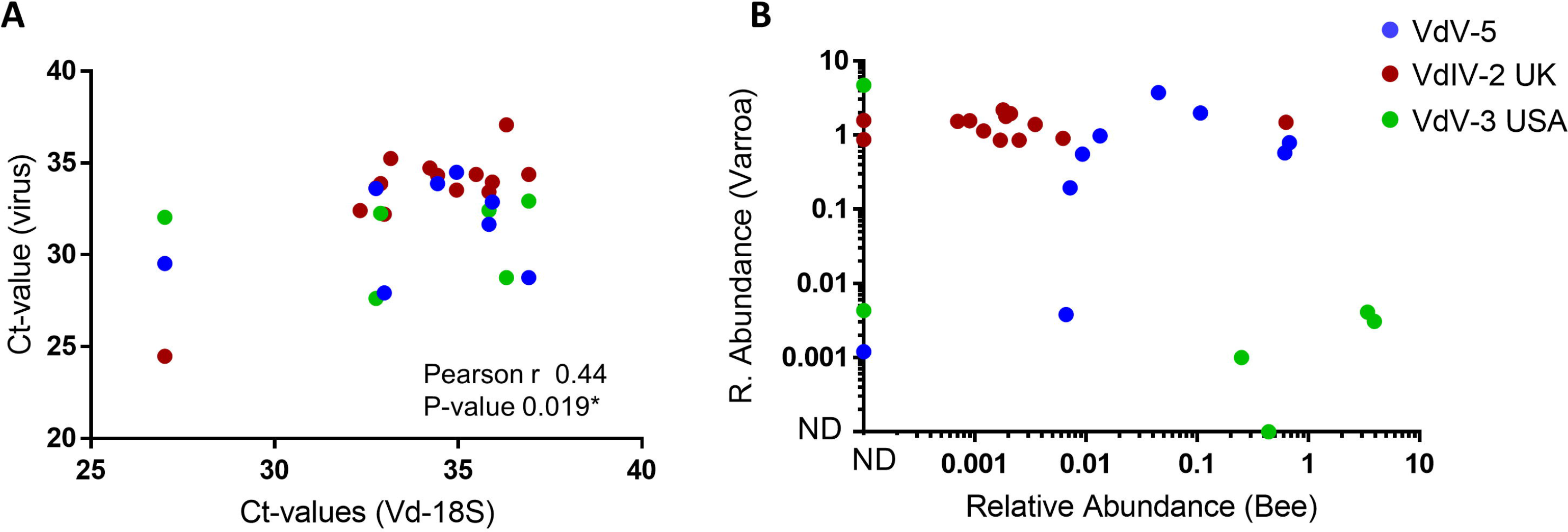
Analysis of the viral abundance in bees. Correlation between the viral abundance (inferred form the Ct-values obtained at the RT-qPCR) in single bees and the amount of Varroa transcripts in the same bee (inferred from the Ct-values of the Varroa reference gene, *18s*) (**A**). Correlation of the relative viral abundance in pairs of immature bees and their parasitizing varroa (**B**). Each dot represents an individual bee (panel **A**) or a honey bee-varroa pair (panel **B**).

Previous evidences of the specific replication of these viruses in *V. destructor* were based on the absence of RT-PCR amplification of viral sequences in the bees (Levin et al. 2016). However, as they used adult worker bees from the colonies to make the detection it is likely that these were not parasitized bees. Hence, there were not exchange of fluids with the mites. In this study, as we used parasitized immature bees, we detected the presence of the three viruses in a large proportion of the analysed bees, but the absence of negative strain clearly indicates that the viruses are not replicating in the bees. Absence of viral replication in the bees does not directly imply the absence of direct or indirect effects on the parasitized bees. For instance, viral particle in the bee tissues could boost or suppress the immunity of the bees. Immunosuppression of honey bee has been associated with the presence of bee-infecting viruses such as DWV (Di Prisco et al. 2016), however direct effects on the bee of non-replicative viruses have not been reported. Alternatively, the replication of these viruses could have an indirect effect interfering or promoting the action of other viruses that replicate actively on the bees.

### Interaction of the new viruses with DWV

The synergistic interaction between *V. destructor* and DWV is one of the major threats to the honey bee industry (Di Prisco et al. 2016). DWV belongs to the *Iflaviridae* family (Valles et al. 2017) so it is also a +ssRNA virus, like the new viruses described here. Therefore, it is possible that its replication and transmission of DWV in the mites could be influenced by the presence of other viruses from the same family (VdIV-2) or with similar replication mechanism (VdV-3 and VdV-5). To test for this potential interference, we determined the abundance of DWV in the studied varroa individuals and the possible correlation with the abundance of the new viruses.

We have a positive detection of DWV in 100 % of the mites tested although with differences in the relative abundance in each case (Fig. 6). To test if these differences in the relative abundance of DWV were influenced by the new viruses, the correlation in their relative abundance were analysed. Viral relative abundances were compared by grouping together the insobeviruses (VDV-3 USA and VDV-5) (Fig. 6A) on one hand and the iflavirus (VdIV-2 UK) on the other (Fig. 6B). No correlation was found in any case (P-values >0.05), so it seems that the newly detected viruses do not interfere with the presence and infection process of DWV and *vice versa*.

**Figure 6.**
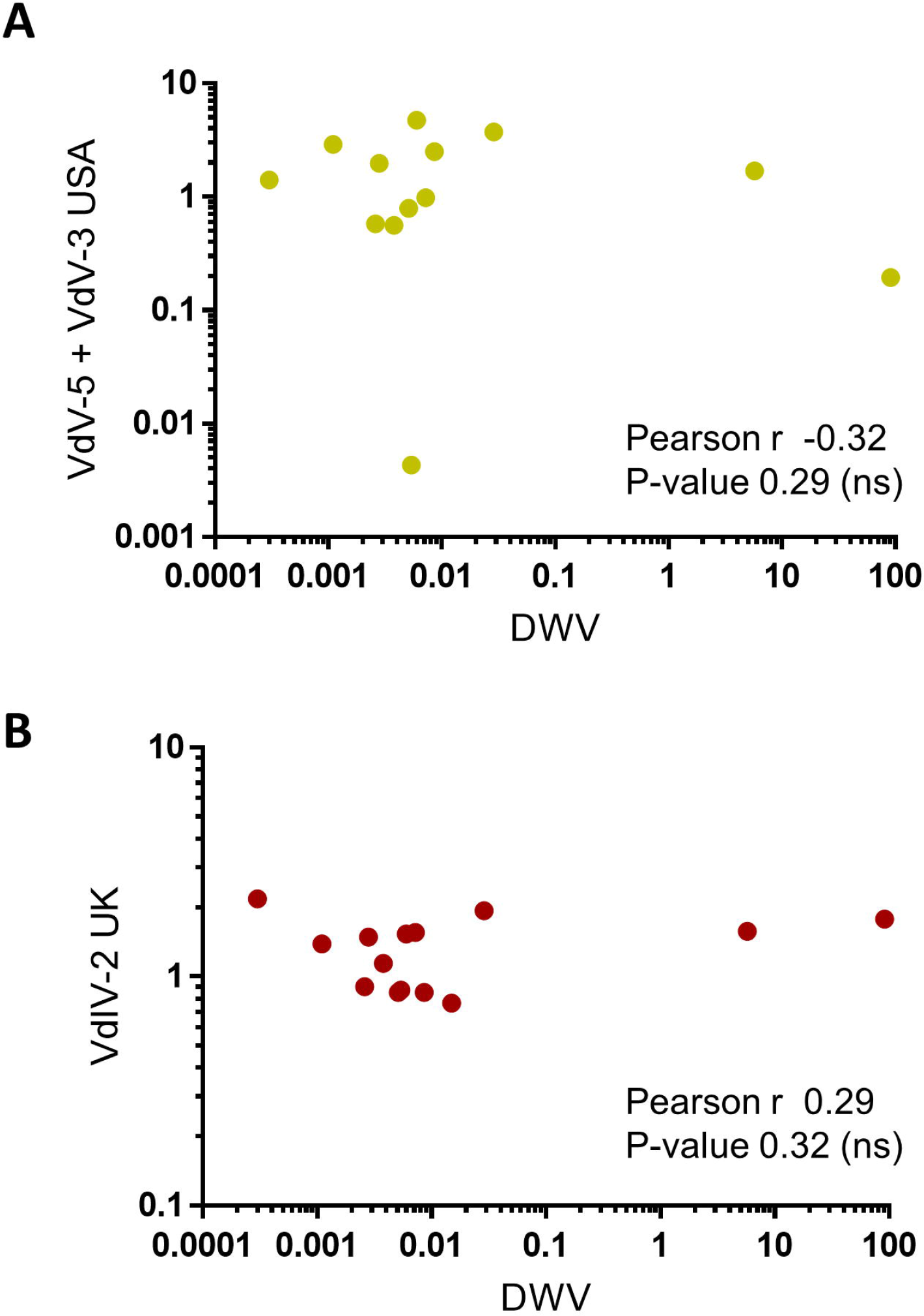
Influence of the novel viruses on DWV abundance on Varroa individuals. Correlation of the relative viral abundance between the VdV-5+VdV-3 USA (**A**) and the VdIV-2 UK (**B**) with DWV in pairs of immature bees and their parasitizing Varroa.

## Conclusion

Three new viruses infecting *V. destructor* have been detected in samples from several locations. We have evidenced that these new viruses actively replicate in *V. destructor* but not in *A. mellifera*, thus to our knowledge this is the first demonstration of Varroa-specific viruses. Although further investigation would be needed to determine the mechanisms of viral transmission and to measure whether they induce a deleterious effect on mite physiology, these new viruses may be used as part of biological control approaches to contribute to the reduction of varroa parasitism on the bee health.

## Acknowledgments

Joel González-Cabrera was supported by the Spanish Ministry of Economy and Competitiveness, Ramón y Cajal Program (RYC-2013-13834). The work at the Universitat de València was funded by a grant from the Spanish Ministry of Economy and Competitiveness (CGL2015‐65025‐R, MINECO/FEDER, UE). Research at the Universitat de València of Stefano Parenti was possible thanks to the Erasmus+ program from the EU. Finally, we thank Fernando Calatayud and Enrique Simó, from the local beekeeper association “apiADS”, for providing bees and mites.

## Supplementary material

**Figure S1. Phylogenetic analysis of the VdIV-2**

Maximum-likelihood phylogenetic tree of the putative RdRp amino acid sequences of *Varroa destructor* iflavirus 2 and closely related members for the *Iflaviridae* and *Dicistroviridae* families. Values on blanches indicates the aLRT support. Branches with aLRT values <◻0.6 have been collapsed. The grey scale bar represents 0.2 amino acid substitution per site. Sequence accession identifiers are provided on Supplementary Table S2.

**Table S1. Sequence of the primers used in the study**

**Table S2. Accession number of viral sequences used in this work for phylogenetic reconstruction**

## References

Boecking O, Genersch E (2008) Varroosis - the ongoing crisis in bee keeping. Journal Fur Verbraucherschutz Und Lebensmittelsicherheit-Journal of Consumer Protection and Food Safety 3, 221–228

Campbell EM, McIntosh CH, Bowman AS (2016) A toolbox for quantitative gene expression in *Varroa destructor*: RNA degradation in field samples and systematic analysis of reference gene stability. PLoS One 11, e0155640

Capella-Gutiérrez S, Silla-Martínez JM, Gabaldón T (2009) trimAl: a tool for automated alignment trimming in large-scale phylogenetic analyses. Bioinformatics (Oxford, England) 25, 1972–3

De Jong D, De Jong PH, Gonçalves LS (1982) Weight loss and other damage to developing worker honeybees from infestation with *Varroa jacobsoni*. J Apicult Res 21, 165–167

Di Prisco G, Annoscia D, Margiotta M, Ferrara R, Varricchio P, Zanni V, Caprio E, Nazzi F, Pennacchio F (2016) A mutualistic symbiosis between a parasitic mite and a pathogenic virus undermines honey bee immunity and health. Proc Natl Acad Sci U S A 113, 3203–8

Di Prisco G, Pennacchio F, Caprio E, Boncristiani HF, Jr., Evans JD, Chen Y (2011) *Varroa destructor* is an effective vector of Israeli acute paralysis virus in the honeybee, *Apis mellifera*. J Gen Virol 92, 151–5

Guindon S, Dufayard J-F, Lefort V, Anisimova M, Hordijk W, Gascuel O (2010) New Algorithms and Methods to Estimate Maximum-Likelihood Phylogenies: Assessing the Performance of PhyML 3.0. Systematic Biology 59, 307–321

Harvey E, Rose K, Eden J-S, Lo N, Abeyasuriya T, Shi M, Doggett SL, Holmes EC (2019) Extensive Diversity of RNA Viruses in Australian Ticks. Journal of Virology 93, e01358–18

Jakubowska AK, Nalcacioglu R, Millan-Leiva A, Sanz-Carbonell A, Muratoglu H, Herrero S, Demirbag Z (2015) In search of pathogens: transcriptome-based identification of viral sequences from the pine processionary moth (*Thaumetopoea pityocampa*). Viruses 7, 456–79

Levin S, Sela N, Chejanovsky N (2016) Two novel viruses associated with the *Apis mellifera* pathogenic mite *Varroa destructor*. Scientific reports 6, 37710

Levin S, Sela N, Erez T, Nestel D, Pettis J, Neumann P, Chejanovsky N (2019) New viruses from the ectoparasite mite *Varroa destructor* Infesting *Apis mellifera* and *Apis cerana*. Viruses 11,

Llopis-Giménez A, María González R, Millán-Leiva A, Catalá M, Llacer E, Urbaneja A, Herrero S (2017) Novel RNA viruses producing simultaneous covert infections in *Ceratitis capitata*. Correlations between viral titers and host fitness, and implications for SIT programs. J Invertebr Pathol 143, 50–60

Longueville J-E, Lefort V, Gascuel O (2017) SMS: Smart Model Selection in PhyML. Molecular Biology and Evolution 34, 2422–2424

McMenamin AJ, Genersch E (2015) Honey bee colony losses and associated viruses. Current Opinion in Insect Science 8, 121–129

Nijveen H, Leunissen JAM, Geurts R, Bisseling T, Rao X, Untergasser A (2007) Primer3Plus, an enhanced web interface to Primer3. Nucleic Acids Research 35, W71–W74

Ongus JR, Peters D, Bonmatin JM, Bengsch E, Vlak JM, van Oers MM (2004) Complete sequence of a picorna-like virus of the genus Iflavirus replicating in the mite *Varroa destructor*. J Gen Virol 85, 3747–55

Pettersson JHO, Shi M, Bohlin J, Eldholm V, Brynildsrud OB, Paulsen KM, Andreassen Å, Holmes EC (2017) Characterizing the virome of Ixodes ricinus ticks from northern Europe. Scientific reports 7, 10870

Potts SG, Roberts SPM, Dean R, Marris G, Brown MA, Jones R, Neumann P, Settele J (2010) Declines of managed honey bees and beekeepers in Europe. J Apicult Res 49, 15–22

Ramsey SD, Ochoa R, Bauchan G, Gulbronson C, Mowery JD, Cohen A, Lim D, Joklik J, Cicero JM, Ellis JD, Hawthorne D, vanEngelsdorp D (2019) *Varroa destructor* feeds primarily on honey bee fat body tissue and not hemolymph. Proc Natl Acad Sci U S A 116, 1792–1801

Roberts JMK, Anderson DL, Durr PA (2017) Absence of deformed wing virus and *Varroa destructor* in Australia provides unique perspectives on honeybee viral landscapes and colony losses. Scientific reports 7, 6925

Rosenkranz P, Aumeier P, Ziegelmann B (2010) Biology and control of *Varroa destructor*. J Invertebr Pathol 103, S96–S119

Sadeghi M, Altan E, Deng X, Barker CM, Fang Y, Coffey LL, Delwart E (2018) Virome of >◻12 thousand Culex mosquitoes from throughout California. Virology 523, 74–88

Santillán-Galicia MT, Ball BV, Clark SJ, Alderson PG (2014) Slow bee paralysis virus and its transmission in honey bee pupae by *Varroa destructor*. J Apicult Res 53, 146–154

Shi M, Lin X-D, Tian J-H, Chen L-J, Chen X, Li C-X, Qin X-C, Li J, Cao J-P, Eden J-S, Buchmann J, Wang W, Xu J, Holmes EC, Zhang Y-Z (2016) Redefining the invertebrate RNA virosphere. Nature 540, 539

Standley DM, Katoh K (2013) MAFFT Multiple Sequence Alignment Software Version 7: Improvements in Performance and Usability. Molecular Biology and Evolution 30, 772–780

Stöver BC, Müller KF (2010) TreeGraph 2: Combining and visualizing evidence from different phylogenetic analyses. BMC Bioinformatics 11, 7

Valles SM, Chen Y, Firth AE, Guérin DMA, Hashimoto Y, Herrero S, De Miranda JR, Ryabov E, Consortium IR (2017) ICTV Virus Taxonomy Profile: Iflaviridae. Journal of General Virology 98, 527–528

Vanengelsdorp D, Evans JD, Saegerman C, Mullin C, Haubruge E, Nguyen BK, Frazier M, Frazier J, Cox-Foster D, Chen Y, Underwood R, Tarpy DR, Pettis JS (2009) Colony collapse disorder: a descriptive study. PLoS One 4, e6481

Webster CL, Longdon B, Lewis SH, Obbard DJ (2016) Twenty-Five New Viruses Associated with the Drosophilidae (Diptera). Evolutionary bioinformatics online 12, 13–25

Yang X, Cox-Foster D (2007) Effects of parasitization by *Varroa destructor* on survivorship and physiological traits of *Apis mellifera* in correlation with viral incidence and microbial challenge. Parasitology 134, 405–412

Yang XL, Cox-Foster DL (2005) Impact of an ectoparasite on the immunity and pathology of an invertebrate: Evidence for host immunosuppression and viral amplification. Proceedings of the National Academy of Sciences of the United States of America 102, 7470–7475

