## Supplementary figures and images for "Identification of new viruses specific to the honey bee mite *Varroa destructor*"

### Supplemental Fig. 1

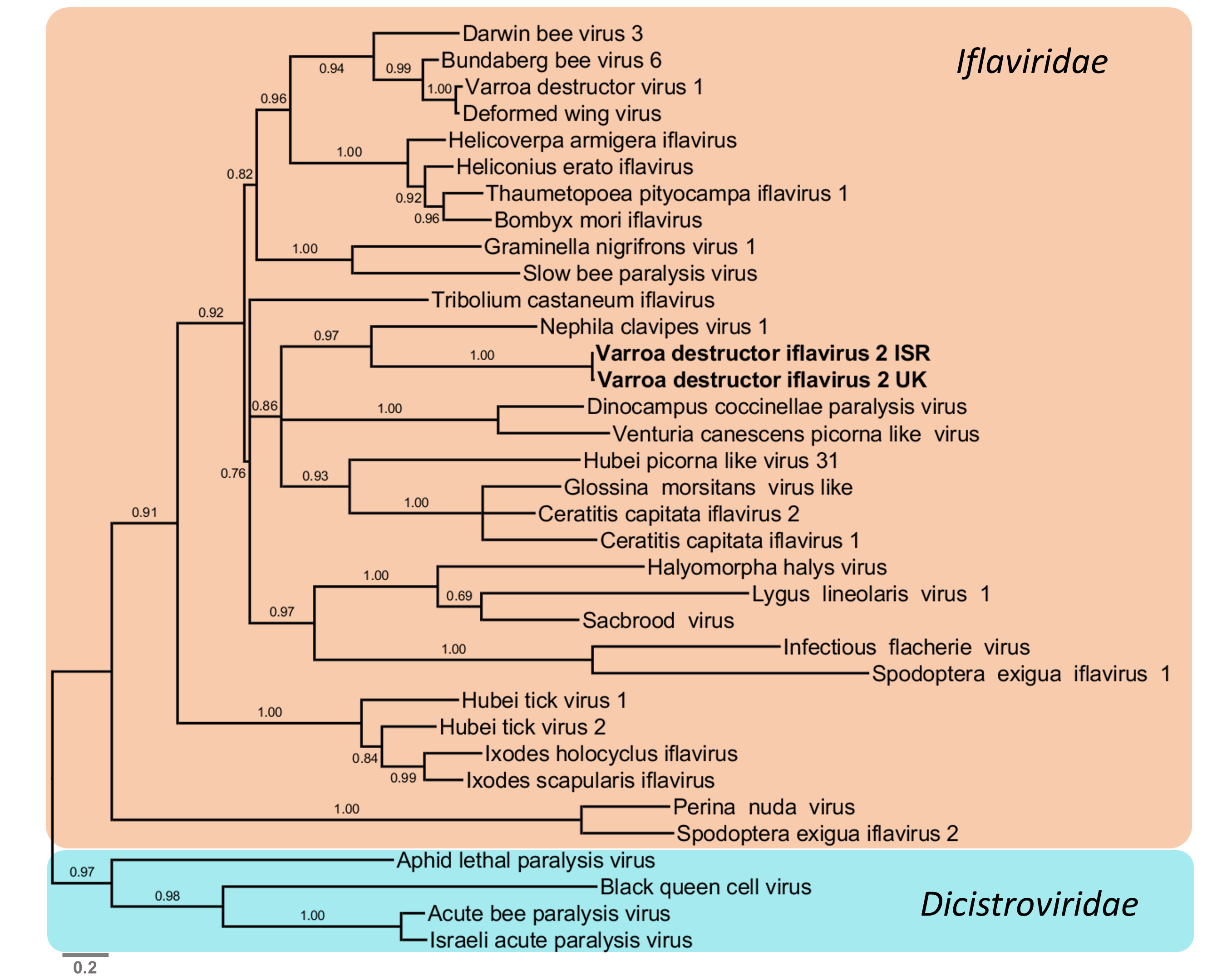
