## Supplemental Table 1 for "Identification of new viruses specific to the honey bee mite *Varroa destructor*"

**Table S1. Sequence of the primers used in the study**

| **Name** | **Sequence (5' - 3')** | **Efficiency (E)** | **Target/application** | **Reference** |
| --- | --- | --- | --- | --- |
| DWV-F | GCGCTTAGTGGAGGAAATGAA | 1.8 | Deform wing virus  RT-qPCR | Di Prisco et al., 2016 |
| DWV-R | GCACCTACGCGATGTAAATCTG |  |  |  |
| VdV-5-F | AGACGTGCTCTTGAGATGGAG | 1.9 | Varroa destructor Virus 5  RT-qPCR | This work |
| VdV-5-R | TCGCGCTTAGCTTCTTTCTC |  |  |  |
| VdV-3 USA F | TCGAACGACCTGAAGAGAAG | 1.8 | Varroa destructor Virus 3 (USA strain). RT-qPCR | This work |
| VdV-3 USA R | GATGGGCATCTGATCATTCC |  |  |  |
| VdIV-2 UK F | ATCCAGATTTGGAGGAGGTG | 1.7 | Varroa destructor Iflavirus 2 (UK strain). RT-qPCR | This work |
| VdIV-2 UK R | TCATCGAGACAATCCTCGTC |  |  |  |
| Vd-18S F | AATGCCATCATTACCATCCT | 1.7 | 18 S Ribosomal RNA gene from *V. destructor.* Endogenous gene for RT-qPCR | Campbell et al., 2016 |
| Vd-18SR | CAAAAACCAATCGGCAATCT |  |  |  |
| ApiDorsal F | TCGGATGGTGCTACGAGCGA | 2.0 | Dorsal gene from *A. mellifera.* Endogenous gene for RT-qPCR | Di Prisco et al 2016 |
| ApiDorsal R | AGCATGCTTCTCAGCTTCTGCCT |  |  |  |
| neg_VdVs F | GGATGCAGGCTACGTGAAGATACG AGATCAATCTGTAATAGATTGAC |  | Forward for cDNA synthesis of negative strand of VdV-5 and VdV-3 USA | This work |
| neg_VdIV2 F | GGATGCAGGCTACGTGAAGATACG CGTCTCCATGCAGAATTAGCTGG |  | Forward for cDNA synthesis of negative strand of VdIV-2 UK | This work |
| TagF | GGATGCAGGCTACGTGAAGATACG |  | Negative strand PCR detection. To be used with VdV-5-R, VdV-3 USA-R, and VdIV-2 UK R | This work |
