## Supplemental Table 2 for "Identification of new viruses specific to the honey bee mite *Varroa destructor*"

**Table S2. Accession number of viral sequences used in this work for phylogenetic reconstruction**

| Name | GenBank Accession | Family |
| --- | --- | --- |
| Acute bee paralysis virus | AF486072.2 | *Dicistroviridae* |
| Aphid lethal paralysis virus | AAN61470.1 | *Dicistroviridae* |
| Black queen cell virus | ABS82427.1 | *Dicistroviridae* |
| Israeli acute paralysis virus | ABY57949.1 | *Dicistroviridae* |
| Bombix mori iflavirus | BAS18834.1 | *Iflaviridae* |
| Bundaberg bee virus 6 | AWK77862.1 | *Iflaviridae* |
| Ceratitis capitata iflavirus 1 | GAMC01001920.1 | *Iflaviridae* |
| Ceratitis capitata iflavirus 2 | GAMC01020602.1 | *Iflaviridae* |
| Darwin bee virus 3 | AWK77848.1 | *Iflaviridae* |
| Deformed wing virus | CAD34006.2 | *Iflaviridae* |
| Dinocampus coccinellae paralysis virus | AIM39350.1 | *Iflaviridae* |
| Glossina morsitans virus-like | ACY69873.1 | *Iflaviridae* |
| Graminella nigrifron virus 1 | AJT58559.1 | *Iflaviridae* |
| Halyomorpha halys virus | AGY34702.1 | *Iflaviridae* |
| Heliconius erato iflavirus | AHW98099.1 | *Iflaviridae* |
| Helicoverpa armigera iflavirus | APW84897.1 | *Iflaviridae* |
| Hubei picorna-like virus 31 | APG77963.1 | *Iflaviridae* |
| Hubei tick virus 1 | APG77503.1 | *Iflaviridae* |
| Hubei tick virus 2 | APG77502.1 | *Iflaviridae* |
| Infectious flacherie virus | BAA25371.1 | *Iflaviridae* |
| Ixodes holocyclus iflavirus | AQZ42314.1 | *Iflaviridae* |
| Ixodes scapularis iflavirus | BBD75427.1 | *Iflaviridae* |
| Lygus lineolaris virus 1 | AEL30247.1 | *Iflaviridae* |
| Nephila clavipes virus 1 | AVK59473.1 | *Iflaviridae* |
| Perina nuda virus | AAL06289.1 | *Iflaviridae* |
| Sacbrood virus | AAD20260.1 | *Iflaviridae* |
| Slow bee paralysis virus | ABS84820.1 | *Iflaviridae* |
| Spodoptera exigua iflavirus 1 | AET36829.1 | *Iflaviridae* |
| Spodoptera exigua iflavirus 2 | AHX00961.1 | *Iflaviridae* |
| Thaumetopoea pityocampa iflavirus 1 | AJC98140.1 | *Iflaviridae* |
| Tribolium castaneum iflavirus | AUE23905.1 | *Iflaviridae* |
| Varroa destructor iflavirus 2 ISR | APB88805.1 | *Iflaviridae* |
| Varroa destructor virus 1 | AAP51418.2 | *Iflaviridae* |
| Venturia canescens picorna like-virus | AAS37668.1 | *Iflaviridae* |
| Mushroom bacilliform virus | AAA53090.1 | *Barnaviridae* |
| Rhizoctonia solani barnavirus 1 | AAA53090.1 | *Barnaviridae* |
| Pea enation mosaic virus 1 | P29154.2 | *Luteoviridae* |
| Poinsettia latent virus | CAI34771.1 | *Solemoviridae* |
| Turnip rosette virus | AAO24320.1 | *Solemoviridae* |
| Cocksfoot mottle virus | ABG73619.1 | *Solemoviridae* |
| Southern bean mosaic virus | ABI53037.2 | *Solemoviridae* |
| American dog tick associated virus 1 | AUX13126.1 |  |
| Baird Spence virus | AOX15243.1 |  |
| Beihai sobemo-like virus 24 | APG75655.1 |  |
| Blue fish point virus | AYP67542.1 |  |
| Culex-associated Luteo-like virus | AXQ04793.1 |  |
| Hubei sobemo-like virus 14 | APG75802.1 |  |
| Hubei sobemo-like virus 16 | APG75906.1 |  |
| Ixodes scapularis associated virus 1 | BBD75429.1 |  |
| Ixodes scapularis associated virus 2 | AII01812.1 |  |
| La Tardoire virus | AMO03215.1 |  |
| Lone star tick associated virus-1 | AUX13123.1 |  |
| Medway virus | AWA82252.1 |  |
| Motts Mill virus | AKH40291.1 |  |
| Norway luteo-like virus 1 | ASY03252.1 |  |
| Norway luteo-like virus 2 | ASY03255.1 |  |
| Norway luteo-like virus 3 | ASY03257.1 |  |
| Norway luteo-like virus 4 | ASY03258.1 |  |
| Shuangao insect virus 9 | APG75749.1 |  |
| Store beach virus | AYP67537.1 |  |
| Tama virus | AWA82270.1 |  |
| Teise virus | AWA82273.1 |  |
| Varroa destructor virus 3 ISR | APB88808.1 |  |
| Wenling sobemo-like virus 1 | APG75959.1 |  |
| Wenzhou sobemo-like virus 3 | APG75759.1 |  |
